# Mouse Models of *GNAO1*-Associated Movement Disorder: Allele- and sex-specific differences in phenotypes

**DOI:** 10.1101/358614

**Authors:** Huijie Feng, Casandra L. Larrivee, Elena Demireva, Huirong Xie, Jeff Leipprandt, Richard R. Neubig

## Abstract

**Background**

Infants and children with dominant *de novo* mutations in *GNAO1* exhibit movement disorders, epilepsy, or both. Children with loss-of-function (LOF) mutations exhibit Epileptiform Encephalopathy 17 (EIEE17). Gain-of-function (GOF) mutations or those with normal function are found in patients with Neurodevelopmental Disorder with Involuntary Movements (NEDIM). There is no animal model with a human mutant *GNAO1* allele.

**Objectives**

Here we develop a mouse model carrying a human *GNAO1* mutation and determine whether clinical features of the *GNAO1* mutation including movement disorder would be evident in the mouse model.

**Methods**

A mouse *Gnao1* knock-in GOF mutation (G203R) was created by CRISPR/Cas9 methods. The resulting offspring and littermate controls were subjected to a battery of behavioral tests. A previously reported GOF mutant mouse knock-in (*Gnao1*^+/G184S^) was also studied for comparison.

**Results**

*Gnao1*^+/G203R^ mutant mice are viable and gain weight comparably to controls. Homozygotes are non-viable. Grip strength was decreased in both males and females. Male *Gnao1*^+/G203R^ mice were strongly affected in movement assays (RotaRod and DigiGait) while females were not. Male *Gnao1*^+/G203R^ mice also showed enhanced seizure propensity in the pentylenetetrazole kindling test. Mice with a G184S GOF knock-in also showed movement-related behavioral phenotypes but females were more strongly affected than males.

**Conclusions**

*Gnao1*^+/G203R^ mice phenocopy children with heterozygous *GNAO1* G203R mutations, showing both movement disorder and a relatively mild epilepsy pattern. This mouse model should be useful in mechanistic and preclinical studies of *GNAO1*-related movement disorders.

## Introduction

Neurodevelopmental Disorder with Involuntary Movements (NEDIM) is a newly defined neurological disorder associated with mutations in *GNAO1*. It is characterized by “hypotonia, delayed psychomotor development, and infantile or childhood onset of hyperkinetic involuntary movements” (OMIM 617493). NEDIM is monogenetic and related to GOF mutations in *GNAO1*, which was previously identified as a gene associated with early infantile epileptic encephalopathy 17 (EIEE17; OMIM 615473).

*GNAO1* encodes Gα_o_, the most abundant membrane protein in the mammalian central nervous system^1^. Gα_o_ is the α-subunit of the G_o_ protein, a member of the G_i/o_ family of heterotrimeric G proteins. G_i/o_ proteins couples to many important G protein-coupled-receptors (GPCRs) involved in movement control like GABA_B_, dopamine D_2_, adenosine A_1_ receptors and adrenergic α_2A_ receptors^2–5^. Upon activation, Gα_o_ and Gβγ separate from each other and modulate separate downstream signaling pathways including Gα_o_ mediated inhibition of cyclic AMP (cAMP), Gβγ mediated inhibition of N-type calcium channels and Gβγ activation of G-protein activated inward rectifying potassium channels (GIRK channels)^6^. G_o_ is expressed mainly in the central nervous system and it regulates neurotransmitter release by modulating intracellular calcium concentrations in pre-synaptic cells^7^. It has also been suggested that G_o_ plays a role in neurodevelopmental processes like neurite outgrowth and axon guidance^8, 9^.

Consequently, G_o_ is an important modulator of neurological functions.

Previously, we defined a functional genotype-phenotype correlation for *GNAO1*^10^: GOF mutations are found in patients with movement disorders, while loss-of-fuction (LOF) mutations are associated with epilepsy^10^. An updated mechanistic review of this genotype-phenotype correlation was recently published^11^. The experimental study of mutant alleles, however, was done with human *GNAO*1 mutations expressed in HET293T cells, which lack a complex physiological content. Therefore, it would be important to see whether mouse models with *GNAO1* mutations would share clinical characteristics of the human patients. Such a result would verify the previously – reported genotype-phenotype correlation and would provide a preclinical testing model for possible new therapeutics. Previously, we reported that heterozygous *Gnao1*^+/G184S^ mice carrying a human-engineered GOF mutation showed heightened sensitization to pentylenetetrazol (PTZ) kindling and had an elevated frequency of interictal epileptiform discharges on EEG^12^. Given the fact that *GNAO1* GOF mutations are associated with movement disorders, we wanted to ask whether *Gnao1*^+/G184S^ mice show both epilepsy and movement disorder phenotypes like 36% of patients with *GNAO1* mutations. G203R is a GOF *GNAO1* mutation that is one of the more common *GNAO1* mutations found clinically^11, 13–17^. Most patients with this mutation exhibit both seizures and movement disorders^11, 13–17^. We wanted to develop a mouse model with that mutation (*Gnao1*^+/G203R^) to see if it replicated the clinical phenotype of *GNAO1* G203R associated neurological disorders.

In this report, we show that two Gα_o_ GOF mutations *Gnao1*^+/G184S^ mice and *Gnao1*^+/G203R^ mice have sex-specific motor impairment and seizures and assess the possibility of using *Gnao1*^+/G184S^ and *Gnao1*^+/G203R^ mice as models to study *GNAO1*-associated neurological defects.

## Materials & Methods

### Animals

Animal studies were performed in accordance with the Guide for the Care and Use of Laboratory Animals established by the National Institutes of Health. All experimental protocols were approved by the Michigan State University Institutional Animal Care and Use Committee. Mice were housed on a 12-h light/dark cycle and had free access to food and water. They were studied between 8-12 weeks old.

### Generation of *Gnao1* mutant mice

*Gnao1*^+/G184S^ mutant mice were generated as previously described^10, 12, 18, 19^ and used as N10 or greater backcross on the C57BL/6J background.

*Gnao1*^G203R^ mutant mice were generated using CRISPR/Cas9 genome editing on the C57BL/6NCrl strain. gRNA targets within exon 6 of the Gnao1 locus (ENSMUSG00000031748) were used to generate the G203R mutation (Figure 1A). Synthetic single-stranded DNA oligonucleotides (ssODN) were used as repair templates carrying the desired mutation and short homology arms (Table 1). CRISPR reagents were delivered as ribonucleoprotein (RNP) complexes. RNPs were assembled in vitro using wild-type S.p. Cas9 Nuclease 3NLS protein, and synthetic tracrRNA and crRNA (Integrated DNA Technologies, Inc.). TracrRNA and crRNA were denatured at 95°C for 5 min and cooled to room temperature in order to form RNA hybrids, which were incubated with Cas9 protein for 5 min at 37°C. RNPs and ssODN templates were electroporated into C57BL/6NCrl zygotes as described previously^20^, using a Genome Editor electroporator (GEB15, BEX CO, LTD). C57BL/6NCrl embryos were implanted into pseudo-pregnant foster dams. Founders were genotyped by PCR (Table 1) followed by T7 Endonuclease I assay (M0302, New England BioLabs) and validated by Sanger sequencing.

**Figure 1.**
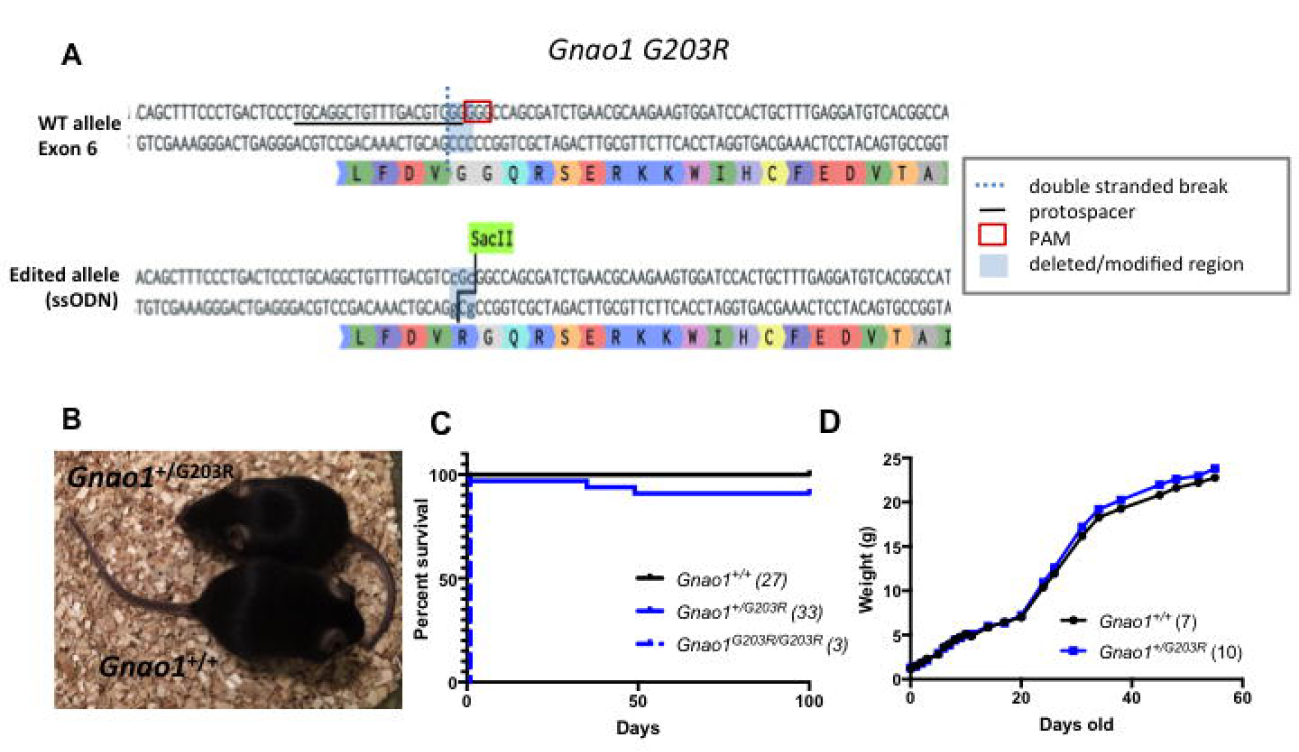
Development of Gnao1^+/G203R^ mouse model. (A) Targeting of the *Gnao1* locus. The location of gRNA target protospacer and PAM, and double stranded breaks following Cas9 cleavage are indicated on the WT allele. Deleted or modified sequences are highlighted in blue. The resulting edited allele sequence and translation are presented along with the sequences used as references for ssODN synthesis. (B) Heterozygous *Gnao1*^+/G203R^ mutant mouse are largely normal in size and behavior. Photo comparing mutant mouse with its littermate control is shown. (C) *Gnao1*^+/G203R^ mice have a relatively normal survival; while homozygous *Gnao1*^G203R/G203R^ mice die perinatally (P0-P1). (D) *Gnao1*^+/G203R^ mice develop normally and gain weight similarly to their WT littermate controls.

**Table 1.**
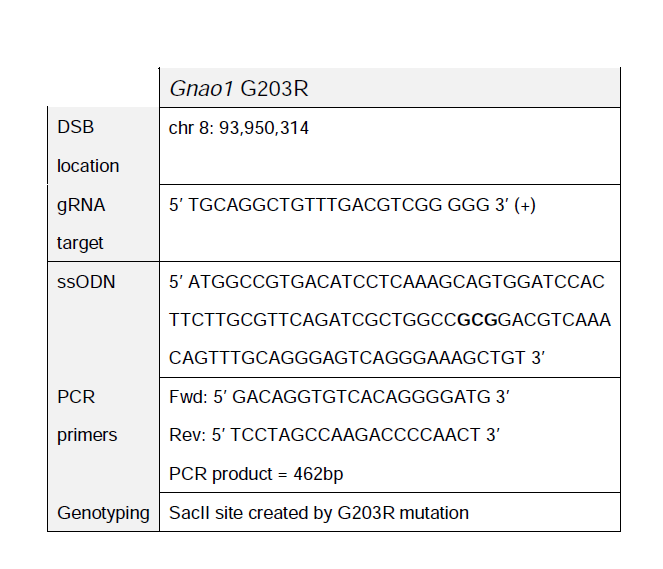
Location, sequence and genotyping of gRNA targets in *Gnao1* locus. gRNA target - 20bp protospacer and PAM sequences are listed, strand orientation indicated by (+) or (-). Sequence of ssODN used as repair template is listed. For G203R, mutated codon is highlighted in bold. DSB - double stranded break. PAM - protospacer adjacent motif.

### Genotyping and Breeding

Heterozygous *Gnao1*^+/G203R^ mutant founder mice were crossed against C57BL/6J mice to generate *Gnao1*^+/G203R^ heterozygotes (N1 backcross). Further breeding was done to produce N2 backcross heterozygotes while male and female N1 heterozygotes were crossed to produce homozygous *Gnao1*^G203R/G203R^ mutants. Studies were done on N1 or N2 G203R heterozygotes with comparisons to littermate controls.

All mice had ears cliped prior before weaning. DNA was extracted from earclips by an alkaline lysis method^21^. The G203R allele of Gα_o_ was identified by Sac II digests (wt 462 Bp and G203R 320 & 140Bp) of genomic PCR products generated with primers (Fwd 5’ GACAGGTGTCACAGGGGATG 3’; Rev 5’ TCCTAGCCAAGACCCCAACT 3’). Reaction conditions were: 0.8µl template, 4µl 5x Promega PCR buffer, 0.4µl 10mM dNTPs, 1µl 10µM Forward Primer, 1µl 10µM Reverse Primer, 0.2µl Promega GoTaq and 12.6 µl DNase free water (Promega catalog # M3005, Madison WI). Samples were denatured for 4 minutes at 95°C then underwent 32 cycles of PCR (95°C for 30 seconds, 60°C for 30 seconds, and 72°C for 30 seconds) followed by a final extension (7 minutes at 72°C). After PCR, samples were incubated with Sac II restriction enzyme for 2 hrs.

### Behavioral Studies

Researchers conducting behavioral experiments were blinded until the data analysis was completed. Before each experiment, mice were acclimated in the testing room for at least 10 min.

### Open Field

The Open Field test was conducted in Fusion VersaMax 42 cm x 42 cm x 30 cm arenas (Omnitech Electronics, Inc., Columbus, OH). Mice and their littermate controls were placed in the arena for 30 minutes to observe spontaneous activities. Using the Fusion Software, distance traveled (cm) was evaluated for novel (first 10 minutes), sustained (10-30 minutes), and total (0-30 minutes) activity. Center Time was also measured. Center Time was defined as the time spent in the center portion (20.32cm x 20.32cm) of the Open Field cage.

### RotaRod

Motor skills were assessed using an Economex accelerating RotaRod (Columbus Instruments, Columbus, OH). The entire training and testing protocol took two days. On day 1, mice were trained for three 2-minute sessions, with a 10-minute rest between each training period. During the first two sessions, the RotaRod was maintained at a constant speed of 5 rpm. In the third training session, the rod was started at 5 rpm and accelerated at 0.1 rpm/sec for 2 minutes. On day 2, mice were trained with two more accelerating sessions for 2 minutes each with a 10-minute break in between. The final test session was 5 minutes long, starting at 5 rpm then accelerating to 35 rpm (0.1 rpm/sec). For all training and test trials, the time to fall off the rod was recorded.

### Grip Strength

Mouse grip strength data was collected following a protocol adapted from Deacon et al^22^ using seven home-made weights (10, 18, 26, 34, 42, 49, 57 grams). Briefly, the mouse was held by the middle/base of the tail and lowered to grasp a weight. A total of three seconds was allowed for the mouse to hold the weight with its forepaws and to lift the weight until it was clear of the bench. Three trials were done to permit the mice to lift the weights with a 10-second rest between each trial. If the mouse successfully held a weight for 3 seconds, the next heavier weight was given; otherwise the maximum time/weight achieved was recorded. A final total score was calculated based on the heaviest weight the mouse was able to lift up and the time that it held it^22^. The final score was normalized to the body weight of each mouse, which was measured before the trial.

### DigiGait

Mouse gait data were collected using a DigiGait Imaging System (Mouse Specifics, Inc., Framingham, MA) ^23^. The test is used for assessment of locomotion as well as the integrity of the cerebellum and muscle tone/equilibrium^24^. Briefly, after acclimation, mice were allowed to walk on a motorized transparent treadmill belt. A high-speed video camera was mounted below to capture the paw prints on the belt. Each paw image was treated as a paw area and its position recorded relative to the belt. Seven speeds (18, 20, 22, 25, 28, 32 and 36 cm/s) were tested per animal with a 5-minute rest between each speed. An average of 4-6 s of video was saved for each mouse, which is sufficient for the analysis of gait behaviors in mice^24^. For each speed, left & right paws were averaged for each animal while fore and hind paws were evaluated separately. Stride length was normalized to animal body length.

### PTZ Kindling Susceptibility

A PTZ kindling protocol was performed as described before^12^ to assess epileptogenesis. Briefly, PTZ (40 mg/kg, i.p. in 5 mg/ml) was administered every other day starting at 8 weeks of age. Mice were monitored and scored for 30 minutes for behavioral signs of seizures as described^12, 25, 26^. Sensitization is defined as death or the onset of a tonic-clonic seizure on two consecutive treatment days. The number of injections for each mouse to reach a sensitization was reported in survival curves.

### Data Analysis

All data was analyzed using GraphPad Prism 7.0 (GraphPad; LaJolla, CA). Data are presented as mean ± SEM and a p value less than 0.05 was considered significant. All statistical tests are detailed in Figure Legends. Multiple comparison correction of the dataset from DigiGait was performed via a false discovery rate (FDR) correction at a threshold value of 0.01 in an R environment using the psych package.

## Results

## *Gnao1*^+/G203R^ mice showed normal viability and growth

Genotypes of offspring of *Gnao1*^+/G203R^ x WT crosses (N1 – C57BL/6NCrl x C57BL/6J) were observed at the expected frequency (29 WT and 27 heterozygous). All three homozygous mice from *Gnao1*^+/G203R^ *x Gnao1*^+/G203R^ crosses died by P1. The small numbers of offspring observed from these crosses so far, however, were not significantly different from expected frequencies (4 wt, 14 het, and 3 homozygous). Heterozygous *Gnao1*^+/G203R^ mice did not show any growth abnormalities compared to *Gnao1*^+/+^ mice (Figure 1B & 1D) and they had relatively normal survival. There were two spontaneous deaths seen for *Gnao1*^+/G203R^ mice out of 33 (Figure 1C). This is reminiscent of the spontaneous deaths seen previously with the *Gnao1*^+/G184S^ GOF mutant mice^12^. *Gnao1*^+/G203R^ mice did not exhibit any obvious spontaneous seizures or abnormal movements.

### Female *Gnao1*^+/G184S^ and male *Gnao1*^+/G203R^ mice show impaired motor coordination and reduced grip strength

Since GOF alleles of *GNAO1* in children result primarily in movement disorder, we tested motor coordination in both our engineered GOF mutant G184S and in the G203R GOF mutant seen in at least 7 children^10, 11^. This was initially assessed using a two-day training and testing procedure on the RotaRod (Figure 2A & B). *Gnao1*^+/G184S^ and *Gnao1*^+/G203R^ mice were compared to their same-sex littermate controls. Female *Gnao1*^+/G184S^ mice exhibited a reduced retention time on the accelerating RotaRod (unpaired t-test, p<0.001, Figure 2A) while male mice remained unaffected. In contrast, male *Gnao1*^+/G203R^ mice exhibited reduced time to stay on a rotating rod (unpaired t-test, p<0.05, Figure 2B) while female *Gnao1*^+/G203R^ mice did not show any abnormalities. Results from all the RotaRod training and testing sessions are shown in Figure S1.

**Figure 2.**
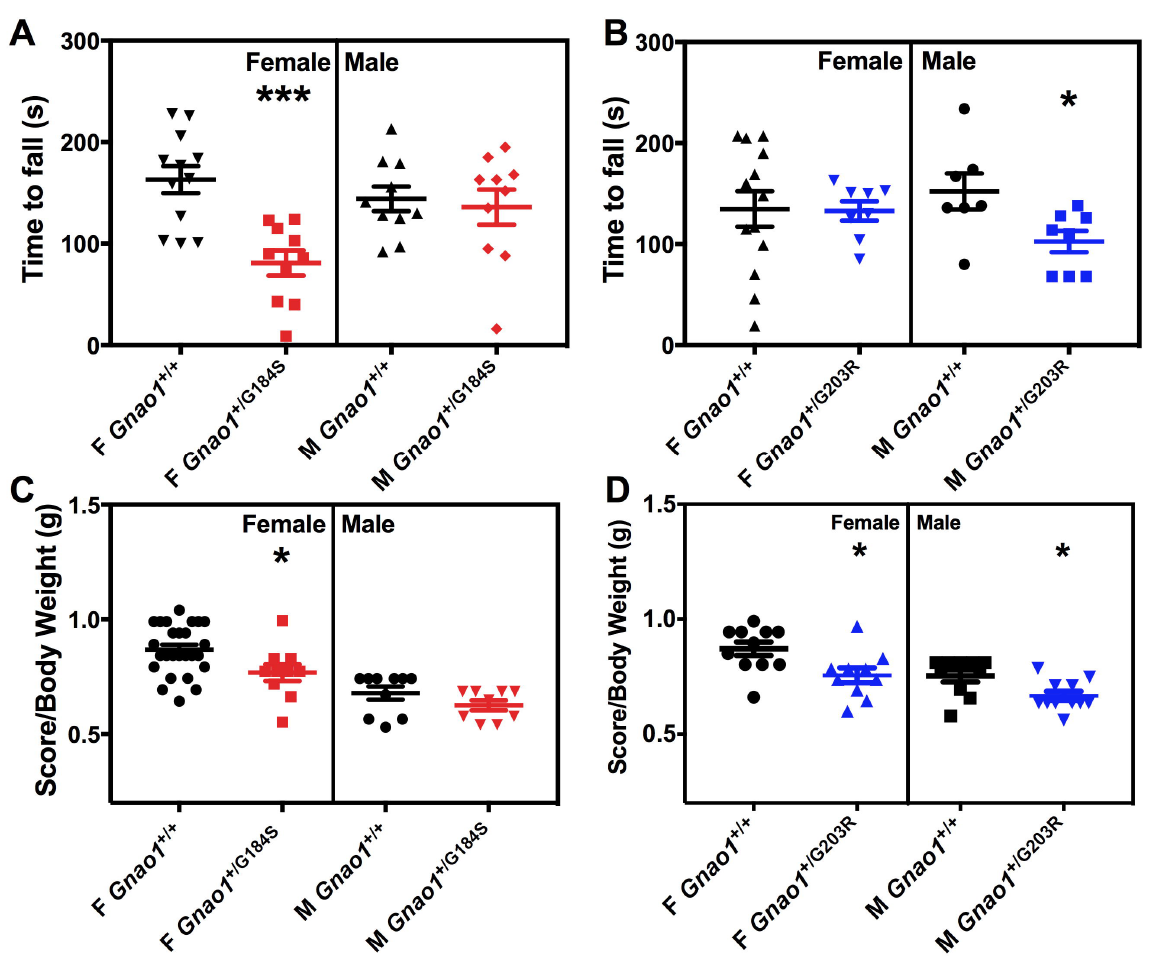
Female *Gnao1*^+/G184S^ mice and male *Gnao1*^+/G203R^ mice show reduced time on RotaRod and reduced grip strength. (A&B) Quantification of RotaRod studies. (A) Female *Gnao1*^+/G184S^ mice lose the ability to stay on a RotaRod (unpaired t-test; ***p<0.001), while male *Gnao1*^+/G184S^ mice appeared unaffected. (B) Male Gnao1^+/G203R also^ showed reduced motor coordination on RotaRod (unpaired t-test, *p<0.01). (C&D) Quantification of grip strength results. Scores for each mouse were normalized to the body weight of the animal measured. (C) Female *Gnao1*^+/G184S^ mice are less capable of lifting weights compared to their *Gnao1*^+/+^ siblings (unpaired t-test, *p<0.05). (D) Both male and female *Gnao1*^+/G203R^ mice showed reduced ability to hold weights (unpaired t-test, *p<0.05). Data are shown as mean ± SEM.

Grip strength was assessed as described ^22^. This test is widely done in combination with the RotaRod motor coordination test. This may be relevant to the hypotonia, seen in many *GNAO1* patients^15, 16, 27–39^. Similar to the RotaRod results, female *Gnao1*^+/G184S^ mice also showed reduced forepaw grip strength compared to their littermate controls (unpaired student t-test, p<0.05, Figure 2C) while males did not exhibit a significant difference (Figure 2C). In contrast, both male and female *Gnao1*^+/G203R^ mice displayed reduced forepaw grip strength (unpaired t-test, p<0.05, Figure 2D).

### *Gnao1*^+/G184S^ mice show reduced activity in the open field arena

The open field test provides simultaneous measurements of locomotion, exploration and surrogates of anxiety. It is a useful tool to assess locomotive impairment in rodents^40^, however, environmental salience may reduce the impact of the motor impairment on behaviror^41^. Therefore, we divided the 30-min open field measurements into two periods, with the first 10 min assessing activity in a novel environment and the 10-30 minute period as sustained activity (Figure 3C & 3D). The novelty measurement showed a significant difference between *Gnao1*^+/G184S^ mice and their littermate controls for both male and female mice (2-way ANOVA, p<0.01 for female, p<0.05 for male, Figure 3C). Female but not male *Gnao1*^+/G184S^ mice showed reduced activity in the sustained phase of open field testing (2-way ANOVA, *p<0.05, **p<0.01, ****p<0.0001). Both male and female *Gnao1*^+/G184S^ mice also showed reduced total activity (2-way ANOVA, p<0.001, Figure 3A & 3C). Neither male nor female *Gnao1*^+/G203R^ mice perform differently in the open field arena compared to their littermate controls (Figure 3B & 3D). No significant difference was observed in the time mice spent in the center of the arena (Figure S2).

**Figure 3.**
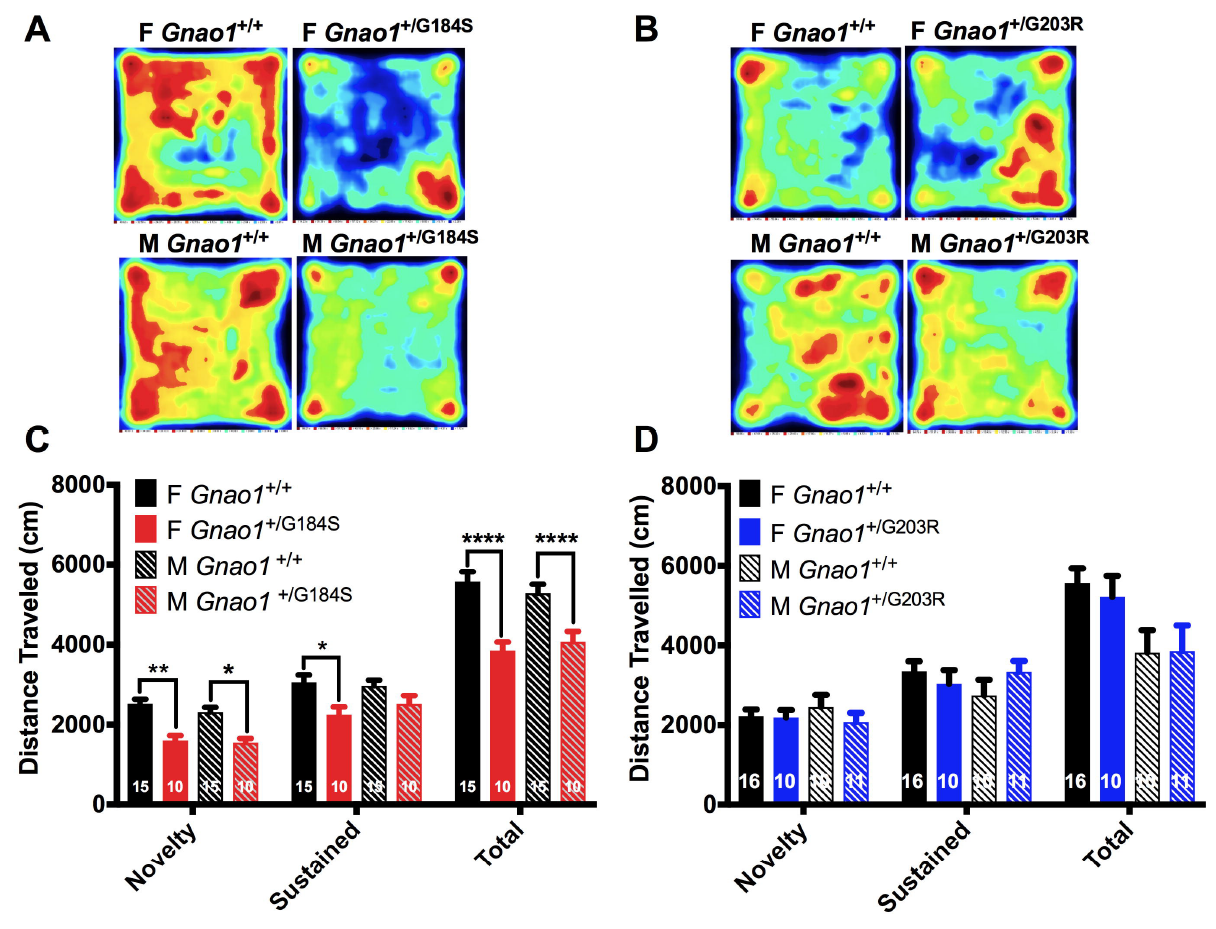
*G184S mutant* mice showed reduced activities in Open Field Test but G203R mutants don’t. (A&C) Female and male *Gnao1*^+/G184S^ mice showed decreased activity in the open field test. A total of 30 min activity was recorded which was divided into Novelty (0-10 min) and Sustained (10-30 min). (A) Representative heat map of overall activity comparison between *Gnao1*^+/+^ and *Gnao1*^+/G184S^ mice in both sexes. (C) Quantitatively, both male and female *Gnao1*^+/G184S^ travelled less in the open field arena (2-way ANOVA; ****p< 0.0001, **p<0.01, *p<0.05). (B & D) Neither male nor female *Gnao1*^+/G203R^ mice showed abnormalities in the open field arena. (B) Sample heat map tracing both female and male mouse movement in open field. (D) Quantification showed no difference between *Gnao1*^+/+^ and *Gnao1*^+/G203R^ mice in distance traveled (cm) in the open field arena (2-way ANOVA; n.s.). Data are shown as mean ± SEM. Numbers of animals are indicated on bars.

### Female *Gnao1*^+/G184S^ mice and male *Gnao1*^+/G203R^ mice exhibit markedly abnormal gaits

In addition to the above behavioral tests, we also performed gait assessment on *Gnao1*^+/G184S^ and *Gnao1*^+/G203R^ mice of both sexes. Gait is frequently perturbed in rodent models of human movement disorders even when the actual movement behavior seen in the animals does not precisely phenocopy the clinical movement pattern^42, 43^. The multiple parameters assessed in DigiGait allow it to pick up subtle neuromotor defects and makes it more informative than the RotaRod test.

The gait analysis largely confirmed the sex differences between the two strains in RotaRod tests. Thirty seven parameters were measured for both front and hind limbs. Given the large number of measurements, we used false discovery rate (FDR) analysis with a Q of 1% as described in Methods to reduce the probability of Type I errors (Figure S4 & S5, Table S1-S4). *Gnao1*^+/G184S^ female mice showed 22 significant differences (Q<0.01) and males showed 8 (Figure S4, Table S3 & S4). For *Gnao1*^+/G203R^ mice, the opposite sex pattern was seen with 27 parameters in females and 8 parameters in males showing significant differences from WT (Figure S5, Table S1 & S2). Two of the most highly significant parameters and ones that had face validity in terms of clinical observations (stride length and paw angle variability) were chosen for further analysis.

Across the range of treadmill speeds, female *Gnao1*^+/G184S^ mice showed significantly reduced stride length (2-way ANOVA, p<0.01, Figure 4A) and increased paw angle variability (2-way ANOVA, p<0.0001, Figure 4C) compared to WT littermates. Male *Gnao1*^+/G184S^ mice only had a difference in paw angle variability ((2-way ANOVA, p<0.0001), not in stride length (Figure 4E & 4G). These results are consistent with the results of RotaRod and grip strength measurements in that female *Gnao1*^+/G184S^ mice showed a stronger phenotype for gait abnormalities than male. In contrast to the *Gnao1*^+/G184S^ mice, male *Gnao1*^+/G203R^ mice appeared to be more severely affected in gait compared to female *Gnao1*^+/G203R^ mice. Male *Gnao1*^+/G203R^ mice had highly significantly reduced stride length (2-way ANOVA, p<0.0001, Figure 4F) and increased paw angle variability (2-way ANOVA, p<0.05, Figure 4H). In contrast, female *Gnao1*^+/G203R^ mice did not show any significant differences in stride length or paw angle variability (Figure 4B & 4D).

**Figure 4.**
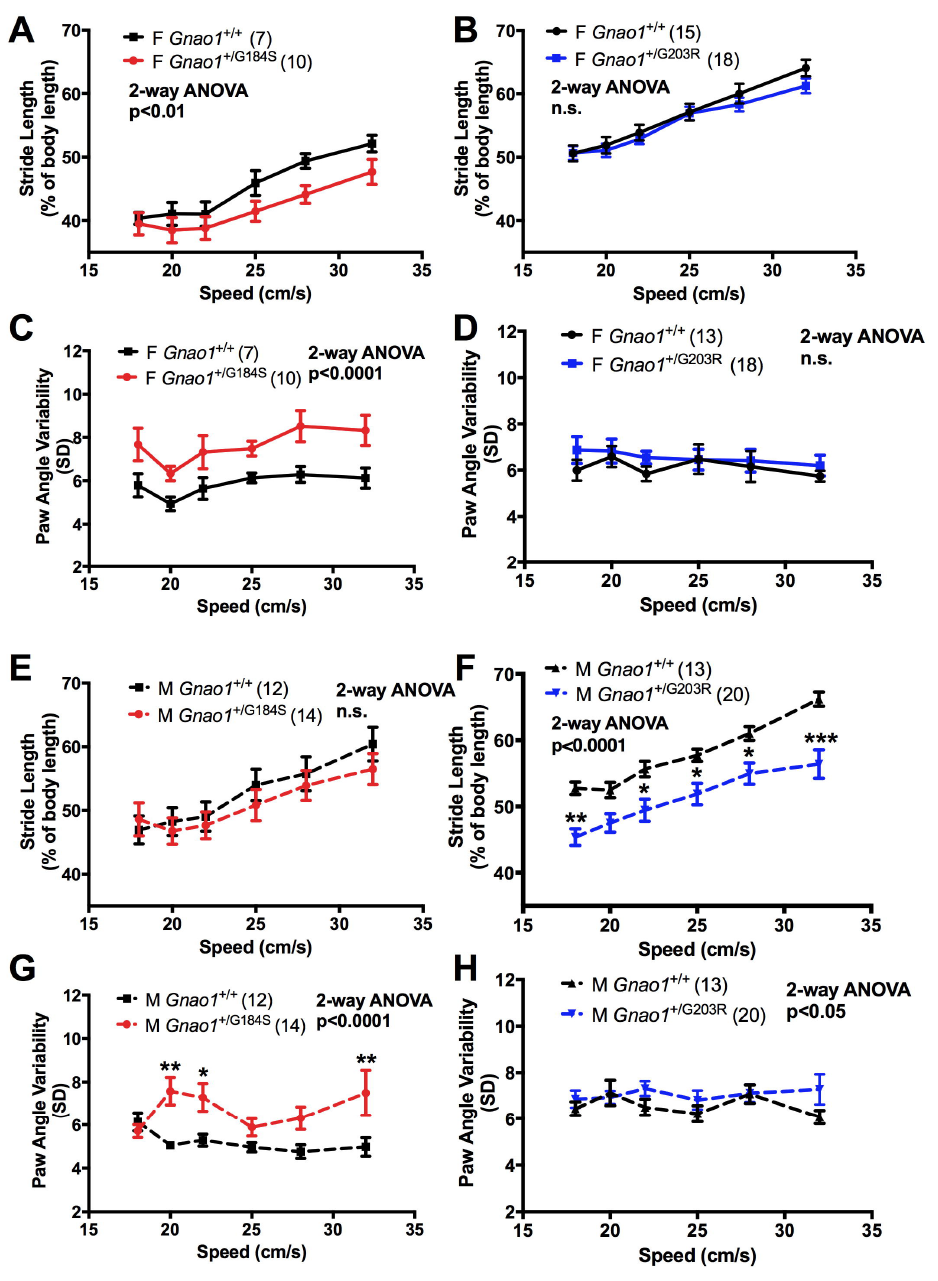
DigiGait Imaging System reveals sex-specific gait abnormalities in *Gnao1*^+/G184S^ mice and *Gnao1*^+/G203R^ mice. (A-D) Female *Gnao1*^+/G184S^ mice showed significant gait abnormalities, while female *Gnao1*^+/G203R^ mice remain normal. (A & B) Female *Gnao1*^+/G184S^ mice showed reduced stride length (2-way ANOVA with Bonferroni multiple comparison post-test) while female *Gnao1*^+/G203R^ mice were unchanged from control (2-way ANOVA; n.s.). (C) Female *Gnao1*^+/G184S^ mice also showed increased paw angle variability (2-way ANOVA, p<0.0001) while female *Gnao1*^+/G203R^ mice showed normal paw angle variability. (E-H) Male *Gnao1*^+/G203R^ and *Gnao1*^+/G184S^ mutant mice showed distinct gait abnormalities. (E & G) Male *Gnao1*^+/G184S^ mice showed significantly increased paw angle variability (2-way ANOVA p <0.0001 overall with significant Bonferroni multiple comparison tests; **p<0.01 and *p<0.05). There was no effect on stride length. (F & H) In contrast, male *Gnao1*^+/G203R^ mice showed markedly reduced stride length (2-way ANOVA p<0.0001 with Bonferroni multiple comparison post-test; ***p<0.001, **p<0.01, and *p<0.05) and modestly elevated paw angle variability (overall p<0.05).

In addition to these quantitative gait abnormalities a qualitative defect was seen. A significant number of *Gnao1*^+/G203R^ mice of both sexes failed to run when the belt speed exceeded 22 cm/s (Mann-Whitney test, female and male p<0.05, Figure 5B). For reasons that are not clear such a difference was not seen for *Gnao1*^+/G184S^ mice (Figure 5A).

**Figure 5.**
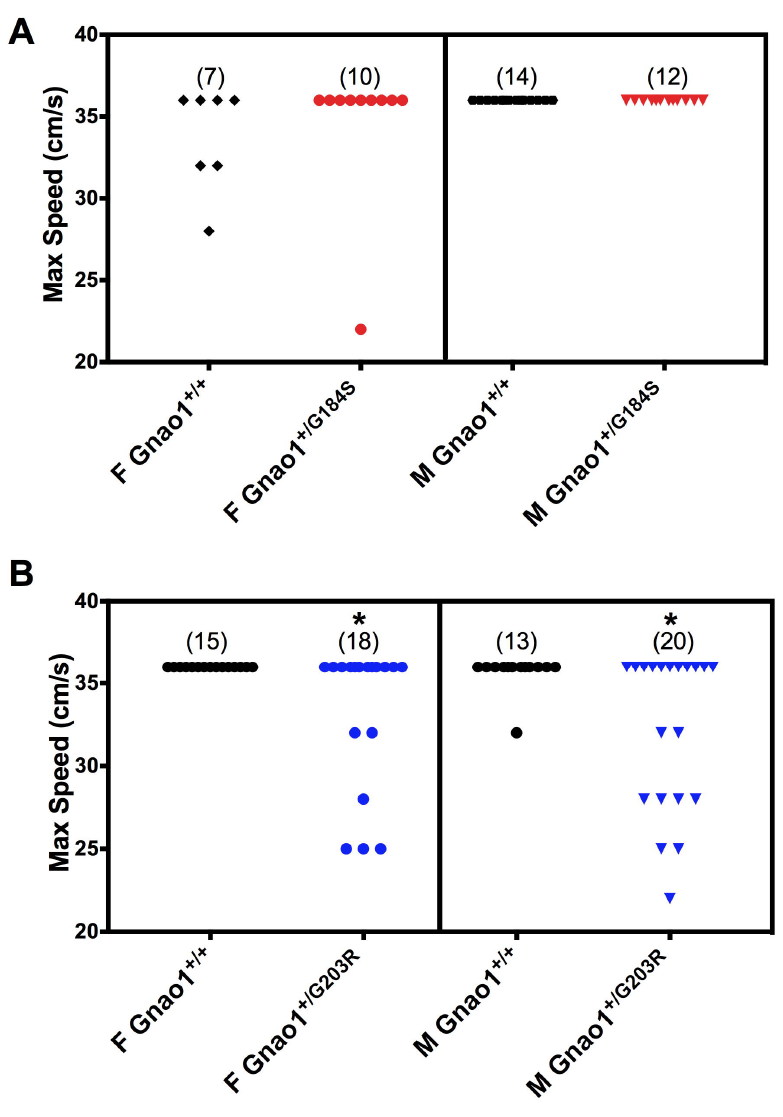
Both male and female *Gnao1*^+/G203R^ mice have reduced maximal running speed on the DigiGait. (A) *Gnao1*^+/G184S^ mice did not show significant differences in the highest treadmill speed successfully achieved. (B) Both male and female *Gnao1*^+/G203R^ mice showed reduced capabilities to run on a treadmill at speeds greater than 25 cm/s (Mann-Whitney test; *p<0.05).

### Male *Gnao1*^+/G203R^ mice are sensitized to PTZ kindling

Epilepsy has been observed in 100% of patients with *GNAO1* G203R mutations^11, 13–15, 17, 44^. Also in the *Gnao1*^+/G184S^ GOF mutant mice, we previously reported spontaneous lethality as well as increased susceptibility to kindling by the chemical anticonvulsant PTZ for both males and females^12^. Kindling is a phenomenon where a sub-convulsive stimulus, when applied repetitively and intermittently, leads to the generation of full-blown convulsions^45^. To determine if the G203R GOF mutant mice mimicked the G184S mutants and phenocopied the human epilepsy pattern of children with the G203R mutation, we assessed PTZ-induced kindling in *Gnao1*^+/G203R^ mutant mice. As expected for C57BL/6 mice, females were more prone to kindling than male mice with half kindled at 4 and 8-10 injections, respectively (Figure 6A & 6B). Despite the increased sensitivity of females in general, female *Gnao1*^+/G203R^ mice did not show significantly higher sensitivity to PTZ compared to their littermate controls (Figure 6A). On the contrary, male *Gnao1*^+/G203R^ mice were more sensitive to PTZ kindling than their controls (Figure 6B, Mantel-Cox Test, p<0.05). Also, three spontaneous deaths were seen (two male and one female) among the 33 G203R mice observed for at least 100 days, similar to the early lethality seen in G184S mutant mice^12^. We cannot, however, attribute those deaths to seizures at this point.

**Figure 6.**
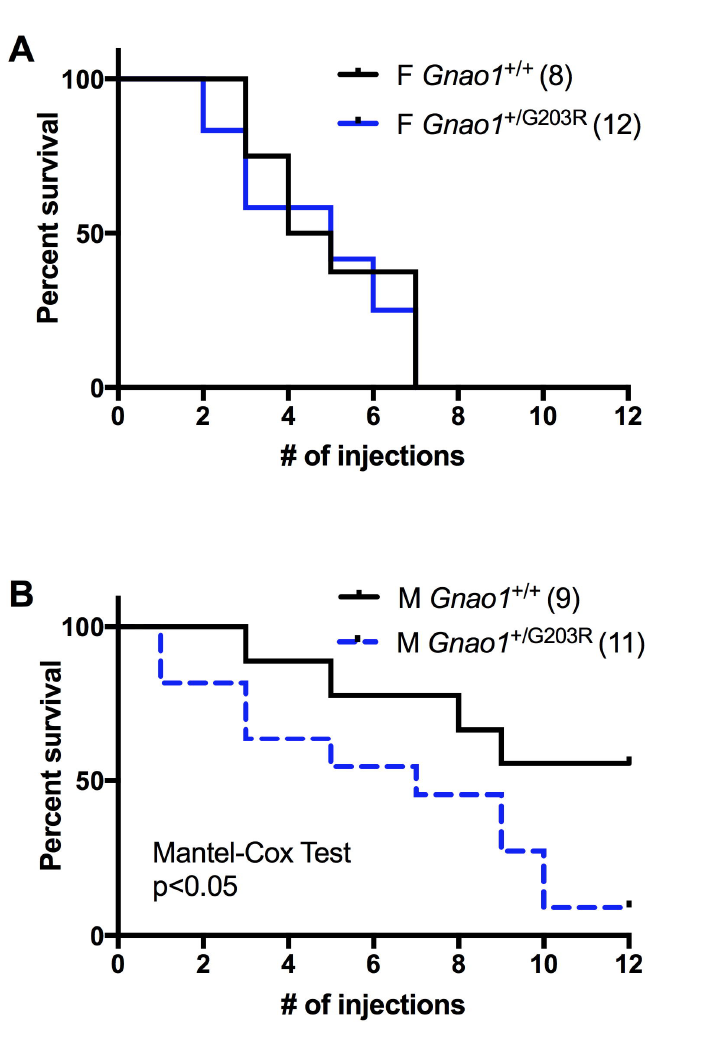
*Gnao1*^+/G203R^ male mice have enhanced Pentylenetetrazol (PTZ) Kindling response. (A) Female *Gnao1*^+/G203R^ did not show heightened sensitivity to PTZ injection. (B) Male *Gnao1*^+/G203R^ mice developed significantly higher sensitivity towards PTZ injections (Mantel-Cox Test; p<0.05).

## Discussion

In this report, we describe the first mouse model carrying a human *GNAO1* mutation associated with disease and we provide evidence to support the concept that GOF mutations are associated with movement disorder^10^. Heterozygous mice carrying the G203R mutation in *Gnao1* exhibit both a mild increase in seizure propensity and evidence of abnormal movements. This fits precisely with the variable seizure pattern of the children who carry this mutation as well as their severe choreoathetotic movements^11, 13–15, 17, 44, 46^. Also in this study, we examined a possible movement phenotype in the experimental mouse GOF mutant (*Gnao1*^+/G184S^) that we reported previously to have a mild seizure phenotype^12^. As predicted from our mechanistic model^10, 11^, it shows movement abnormalities as well.

In mouse models of movement disorders, the mouse phenotype is not as striking or as easily observed as the clinical abnormalities in the patients^47, 48^, however they are often informative about mechanism and therapeutics. For the patient-derived *Gnao1*^+/G203R^ mutant mouse, neither the seizure propensity nor the movement abnormality was obvious without a stress being applied. Male *Gnao1*^+/G203R^ mice showed decreased motor ability on RotaRod, decreased fore paw strength, and gait abnormalities at higher speeds of walking/running. No spontaneous seizures were observed but there was a substantial increase in sensitivity to PTZ-induced seizures in the kindling model. This very closely replicates the mild seizure phenotype of *Gnao1*^+/G184S^ mice^12^. We now show that the female *Gnao1*^+/G184S^ mice also exhibit gait and motor abnormalities.

Both the *GNAO1* G203R and the G184S mutations show a definite but modest GOF phenotype in biochemical measurements of cAMP regulation^10^. In each case, the maximum percent inhibition of cAMP is not greatly increased but the potency of the α_2a_ adrenergic agonist, used in those studies to reduce cAMP levels, was increased about 2-fold. This effectively doubles signaling through these two mutant G proteins at low neurotransmitter concentrations (i.e. those generally produced during physiological signaling). This, however, does not prove that cAMP is the primary signal mechanism involved in pathogenesis of the disease. The heterotrimeric G protein, G_o_, of which the *GNAO1* gene product, Gα_o_, is the defining subunit, can signal to many different effector mechanisms^8, 11, 49^. We recently reviewed the mutations associated with genetic movement disorders and identified both cAMP regulation and control of neurotransmitter release as two *GNAO1* mechanisms that seem highly likely to account for the pathophysiology of *GNAO1* mutants^11^. Since many G_o_ signaling effectors (including cAMP and neurotransmitter release) can be mediated by the Gαγ subunit released from the G_o_ heterotrimer, other effectors could also be involved in the disease mechanisms. A recent hypothesis has also been raised that intracellular signaling by Gα_o_ may be involved^50^. The observation that one of the most common movement disorder-associated alleles (R209H and other mutations in Arg^209^) does not markedly alter cAMP signaling in *in vitro* models, does suggest that the mechanism is more complex than a simple GOF vs LOF distinction at cAMP regulation.

We observed a striking sex difference in the phenotypes of our two mouse models. Female *Gnao1*^+/G184S^ mice and male *Gnao1*^+/G203R^ mice showed much more prominent movement abnormalities than male G184S and female G203R mutants. However, the patterns of changes in the behavioral tests did not exactly overlap. Only G184S mutants showed significant changes in open field tests while only the G203R mutants showed the striking reduction in ability to walk/run at higher treadmill speeds. For both mutant alleles, the seizure phenotype was also worse in the sex with more prominent movement disorder. *GNAO1* encephalopathy is slightly more prevalent (60:40) in female than male patients^11^. It is not uncommon to have sex differences in epilepsy or movement disease progression. One possible explanation is that estrogen prevents dopaminergic neuron depletion by decreasing the uptake of toxins into dopaminergic neurons in Parkinson’s disease (PD) animal model induced by neurotoxin^51^. The G_i/o_ coupled estrogen receptor, GPR30, also contributes to estrogen physiology and pathophysiology^52^. PD is more common in male than female human patients^53^, therefore, the pro-dopaminergic properties of estrogen may exacerbate conditions mediated by hyper-dopaminergic symptoms like chorea in Hungtington’s disease (HD)^51^. Chorea/athetosis is the most prevalent movement pattern seen in *GNAO1*-associated movement disorders^11^ so the female predominance correlates with that in HD. Clearly mechanisms of sex differences are complex including differences in synaptic patterns, neuronal densities and hormone secretion^51, 54, 55^, but it is beyond the scope of this report to explain how the molecular differences contribute to the distinct behavioral patterns.

Since *GNAO1* encephalopathy is often associated with developmental delay and cognitive impairment^11^, it would be interesting to see whether the movement phenotype we have seen in female *Gnao1*^+/G184S^ and male *Gnao1*^+/G203R^ mice is due to a neurodevelopmental malfunction or to ongoing active signaling alterations. G_o_ coupled GPCRs play an important role in hippocampal memory formation^56, 57^. Additional behavioral tests will be valuble to assess the learning and memory ability of the *Gnao1* mutant mice.

With the increasing recognition of *GNAO1*-associated neurological disorders, it is important to learn about the role of G_o_ in the regulation of central nervous system. The novel *Gnao1* G203R mutant mouse model reported here, and further models under development, should facilitate our understanding of *GNAO1* mechanisms in the *in vivo* physiological background rather simply in *in vitro* cell studies. The animal models can also be used for preclinical drug testing and may permit a true allele-specific personalized medicine approach in drug repurposing for the associated movement disorders.

## Acknowledgements

This work was supported by a pre-doctoral fellowship from the American Epilepsy Society (H.F) and funds from the Michigan State University Clinical and Translational Science Institute.

## Authors’ Role

1) Research project: A. Conception: H. Feng, R. R. Neubig; B. Organization: H. Feng, C. L. Larrivee, R. R. Neubig; C. Execution: E. Demireva, H. Xie, H. Feng, C. L. Larrivee, J. Leipprandt

2) Statistical Analysis: A. Design: H. Feng, C. L. Larrivee, R. R. Neubig; B. Execution: H. Feng, C. L. Larrivee, R. R. Neubig; C. Review and Critique: R. R. Neubig

3) Manuscript: A. Writing of the first draft: H. Feng; B. Review and Critique: R. R. Neubig, C. L. Larrivee

